# Enriched environmental intervention mitigates hippocampal cellularity and behavioral disorders in maternal protein-restricted male rat offspring

**DOI:** 10.1101/2024.05.30.596620

**Authors:** Gabriel Boer Grigoletti-Lima, Patrícia Aline Boer, José Antonio Rocha Gontijo

## Abstract

**Background:** Gestational protein intake restriction induces long-lasting harmful outcomes in the offspring’s organs and systems.

**Aims:** This study sought to evaluate the effects of protein restriction during pregnancy and breastfeeding in 42-day-old male offspring on the structure of the hippocampus, behavior tests related to memory and emotions, and the influence of an enriched environment on these parameters.

**Results and Discussion:** The current study demonstrated that maternal protein restriction during neural development causes crucial morphological changes in the hippocampus, making the LP offspring vulnerable to specific neural disorders in adulthood. In addition, it supports the ‘selfish brain’ theory, a paradigm that postulates the brain maintains its mass ‘selfishly’ by reallocating resources from other body parts when faced with nutritional stress. However, the hippocampus cellularity pattern was profoundly altered, significantly reducing the number of neurons after the breastfeeding period. This may expand the understanding of nutritional stress affecting the brain area’s constitution and its supposed effects on posterior behavioral disorders. Here, reciprocal data was observed between brain masses, changes in the hippocampus cell pattern, and decreased body mass in the LP progeny.

**In conclusion** it was demonstrated that neuronal composition and structure profoundly modified by dietary restriction are surprisingly restored from primordial cells by exposure to the enriched environment. In addition, we must emphasize that although we have observed a significant reduction in the number of neurons after gestational and breastfeeding periods, we demonstrated for the first time a substantial reduction in the fear-reflecting behavior, which an enriched environment exposure may revert. The enriched environment also significantly modified the discrimination ratio, increasing the ability of both progenies to discriminate between novel and familiar objects in a short time associated with reverse abnormal hippocampus cell patterns. These findings underscore the potential for environmental interventions to mitigate the effects of early-life nutritional stress on brain development and behavior.

## 1. INTRODUCTION

The rodent’s morphological and functional organization of the brain regions is established during the prenatal and postnatal periods, involving proliferation and differentiation of neural cellular components, non-neuron onset, and cell migration. Several environmental factors, such as maternal nutrition and environmental insults, affect all these processes, which could lead to changes in the long-term offspring’s brain morphology and functionality [1–5]. Maternal undernutrition is an essential factor determining epigenetic modifications in offspring’s intrauterine central nervous system (CNS) development, leading to permanent disorders, including learning and memory deficits [6–8]. Children exposed to malnutrition in perinatal periods showed cognitive deficits [9,10] and a high risk of developing psychiatric disorders, such as depression [11,12] and schizophrenia [13,14]. Studies in animal models submitted to gestational malnutrition have shown that adult offspring have significant learning and memory deficits [15–17]. Besides, these animals exhibit an excessive response to stressors [18,19] and a predisposition to adding psychotropic drugs [20]. However, cellular and molecular mechanisms of these neurobehavioral disorders are not entirely known in maternal protein-restricted offspring.

The hippocampus is a brain structure related to emotion, memory, and spatial learning. Its structure is divided into the ventral hippocampus (VH) and the dorsal hippocampus (HD) [21]. In information processing, the HD is responsible for spatial memory, while the HV responds to stress and emotions [21,22]. Histologically, the hippocampus is divided into subfields CA1, CA3, and dentate gyrus (DG), and the subventricular zone (SVZ) is a region of many stem cells, even in adulthood, that migrates to the hippocampus subfield and olfactory bulb and differentiate into local interneurons [23].

Previous studies have demonstrated that severe maternal protein restriction (6%) in the rat negatively programs the time course of offspring development of the DG, [24,25] the morphology of hippocampal cells and synaptic spine complexity26 and the number and distribution of neurotransmitter receptors [8,27,28].

Reyes-Castro et al. [29], using a model of 10% protein restriction in both gestational and lactation periods, showed, on postnatal day 110, impairment of spatial acquisition and memory retention in male offspring, alterations in hippocampal presynaptic and postsynaptic (spines) elements, and higher glucocorticoid and ACTH levels. The authors suggest these are potential mechanisms to explain programmed rats’ learning and memory deficits.

Conversely, the enriched environment (EE) comprises complex inanimate and social stimuli that benefit physiological processes, such as improving learning and memory and reducing cognitive decline associated with aging [30]. The enriched environment also contributes to increased hippocampus synaptogenesis, dendritic spines, and synapses in some neuronal populations [31,32]. In the DG, the EE increases neurogenesis and reduces cell death by apoptosis [33–35]. So, several studies established that EE promotes changes in the brain at molecular, anatomical, and functional levels [30,33,36–39]. Thus, gestational and prenatal malnutrition may have more harmful effects on the neural development of rats. The maturation and development of the central nervous system also depend on environmental stimulation and enrichment. Therefore, it hypothesized that gestational and lactation protein restriction would change the offspring’s hippocampi neurons and non-neuron pattern and whether an enriched environment may repair this change.

The hippocampus was the focus because this structure is intimately involved in learning and memory and is particularly susceptible to developmental disturbances and the progenitor cells niche within the DG. So, this study aimed to determine the effects of maternal protein restriction on hippocampal cell patterns and neural stem cells associated with changes in learning, memory, and behavioral tests in offspring compared to an appropriate control group.

## 2. MATERIAL AND METHODS

### 2.1. Animals and experimental groups

Experiments were conducted on age-matched sibling-mated Wistar HanUnib rats (250–300g). General guidelines established by the Brazilian College of Animal Experimentation (COBEA) and approved by the Institutional Ethics Committee (CEUA/UNICAMP #3634-1) were followed throughout the study. Our local colonies originated from breeding stock supplied by CEMIB/UNICAMP, Campinas, SP, Brazil. Immediately after weaning at three weeks of age, animals were maintained under controlled temperature (25°C) and lighting conditions (08:00–18:00 h), with *ad libitum* access to tap water and standard rodent laboratory chow (Nuvital, São Paulo, Brazil) rats were raised to 12 weeks of age. Dams were maintained in an isocaloric standard rodent laboratory with *ad libitum* access to either regular protein content (NP) (17% protein, n = 40) or low protein content (LP) (6% protein, n = 40) chow (Table 1) for the duration of their pregnancy. When rats were placed to mate, sperm in the vaginal smear was designated as day 1 of pregnancy. Dams were kept in individual metabolic cages and weighed during pregnancy. On the birthday, male offspring were weighed; keeping eight pups per dam reduced the litter, and only male offspring were used. During breastfeeding, the mothers continued to be fed with NP and LP diet. After weaning (21° day), the offspring were fed with a standard rat diet, and the following groups were formed: (1) NP rats submitted to the standard environment (NPS, n = 20); (2) NP rats submitted to the enriched environment (NPE, n = 20); (3) rats from the LP group submitted to the standard environment (LPS, n = 20) and (4) rats from the LP group submitted to the enriched environment (LPE, n = 20). The enriched environment (EE) includes colored pet cages with several tubes, ramps, and exercise wheels. Colored wood objects were offered for exploration. After 21 days in the standard or enriched environment, the animals were submitted to behavioral tests described below.

**Table 1.**
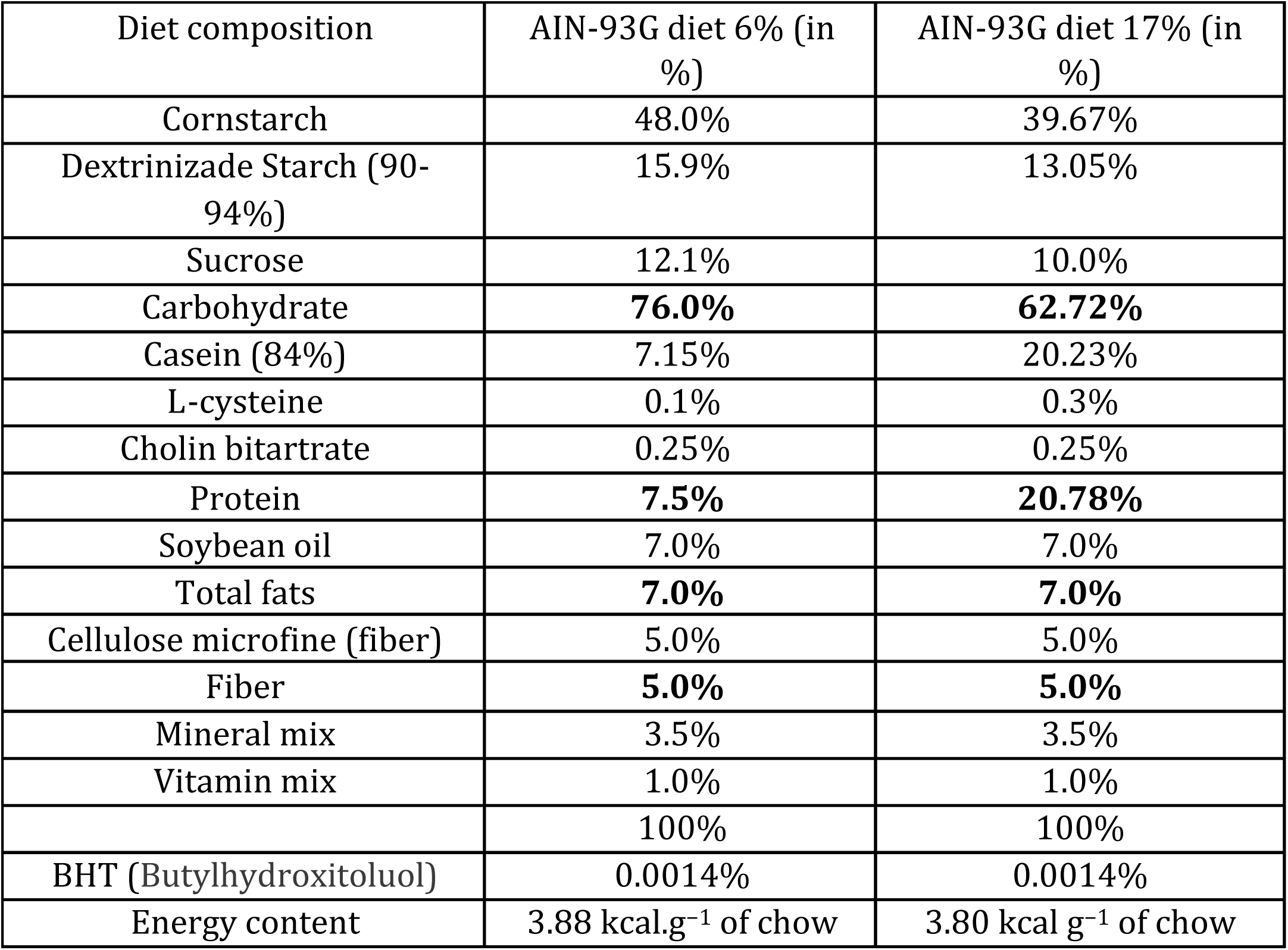
Composition of standard rodent laboratory diet (standard normal-protein (NP) diet, 17%, and low-protein (LP) diet, 6% (AIN 93G). This diet composition played a crucial role in our study, influencing the behavior of the rats.

### 2.2 Behavioral Testing

Beyond 42 days after birth, the male offspring from each litter, NPE and NPS (n=20 for each group), LPE, and LPS (n=20 for each group), were used for behavioral research. All behavioral tests were performed during the light cycle. Test room illumination was kept constant and controlled under low-intensity white lighting (5–30 lux).

#### 2.2.1 Activity Monitoring Test

42-days-old LP and NP offspring subgroups, in a standard or enriched environment, were tested for 5 min in an activity monitor (EP149 Insight, Brazil) equipped with six bars with 16 infrared sensors that detect and monitor the relative position of the animal in the acrylic box (500 x 480 x 500 mm). The movements are captured in the x, y, and z-axes. The program detected circular, time, and counterclockwise movements, walking and resting time, ambulatory and unstable (or stereotyped) movements, rotational movements, number of jumps, and time spent in an orthostatic position and still can measure the space traveled and time. The animals were placed in the center of the monitor box, and their movements were recorded for 5 minutes. Also, at the end of each experimental session, the % of fecal cakes (defined as evidence of fear) was checked [40]. Then, the equipment was cleaned with 5% v / v ethyl alcohol to remove the previous animal smell.

#### 2.2.2 Elevated plus maze (EPM)

42-days-old LP and NP offspring subgroups were tested in a standard or enriched environment for 5 min in an EPM with Software Monitoring Sensors (EP151MO Insight, Brazil). The animals were placed in the center arena with their heads facing the closed space, and the duration of time spent in the open arms versus the closed arms was used as an anxiety index. The percentage of open arm entries (number of open arm entries/total entries) x100, the percentage of open arm time (open arm time/total test time) x100, and the duration of time spent in the open arms versus the closed arms were used as an anxiety index. In contrast, the total number of arm entries and the number of closed arm entries also served as measures of general motor activity.

#### 2.2.3 Object Discrimination Test

After adapting the animals to the experimental environment in acrylic boxes, the animals of the different groups were placed in an environment with similar objects for 5 minutes. To study short-term memory, 3 hours after the first exposure, the animals were placed in front of the objects. However, one was exchanged for another of a different color and shape for 5 minutes. The entire test was filmed. After 24 hours, the long-term memory study was carried out when one of the initially exposed objects was exchanged for another one with a different color and shape, subjecting the animals to exposure to this situation for 5 minutes as well. The animals were considered to explore the objects when they approached the muzzle by approximately 1 cm from the objects. The exploration time (t) of each object was noted. The discrimination ratio (D) was calculated as the ratio between the exploration time of the new object subtracted from the exploration time of the familiar object divided by the total exploration time of both the objects expressed in percentage, as follows the formula: D=(t [new]-t [familiar]/t [new]+t [familiar]) x100.

#### 2.2.4. Morris water maze test (MWM)

was performed as previously described [6]. Briefly, the tests were conducted in a circular black tank (170 cm diameter with a depth of 31 cm, at 22 ◦C) in a dimly lit room with clues in the walls. The tank was divided into four imaginary quadrants with a platform (12 cm diameter, 30 cm height) in one of them. After working and reference memory tests, the escape platform is removed on the eighth day, and the rodents can search for it for 60 seconds. The working memory task is a test of PFC function, and its goal is to assess the ability of rats to learn the position of the hidden platform during four consecutive trials. It consisted of 4 days of acquisition (4 trials/day). The platform’s position was kept constant on each trial day, but the position varied on each successive day. A trial was considered to have ended when the rat escaped onto the platform. The time spent to reach the platform (escape latency) was recorded. The reference memory task assesses the function of the hippocampus, and its goal is to determine the ability of rats to learn the position of the hidden platform and to keep this information during all test days. The test consisted of 4 days of acquisition (4 trials/day). During the four days, the platform remained in the same quadrant. A trial was considered to have ended when the rat escaped onto the platform. The time (in seconds) spent to reach the platform (escape latency) was recorded in the consecutive trials. To assess reference memory and retention at the end of learning, 24 h after training on the eighth day, rats were returned to the water maze for a probe trial. The hidden platform was removed, and mice were allowed to swim freely for 60 seconds. The time spent in the quadrant where the platform was previously located relative to the other three quadrants was used to index the mice’s memory capacity.

### 2.3 Hippocampal total cell and neuron quantification

The cell and neuron quantification followed the technique previously described [17, 41]. Five 42-day-old offspring from different mothers of NPS, NPE, LPS, and LPE rats were sacrificed and perfused transcardially with saline, followed by 4% buffered paraformaldehyde. The isolated hippocampus was mechanically dissociated and homogenized in a solution of 40 mM sodium citrate and 1% Triton X-100, as previously described [40,41]. After several wash and centrifugations, nuclei were suspended in a solution (phosphate-buffered saline containing 1% 4’, 6-diamidino-2-phenylindole (DAPI) (Molecular Probes, Eugene, OR, USA), and nuclei density was determined in a hemocytometer. The total number of cells was estimated by multiplying the mean nuclear density by the total suspension volume. For estimates of total neuron number, a 200–500-µl aliquot is removed from the nuclear suspension and incubated in an anti-NeuN antibody (1:300 in PBS; Chemicon, Temecula, CA, USA). After washing, nuclei are incubated in Cy3-conjugated secondary antibody (1:400 in 40% PBS, 10% goat serum, and 50% DAPI; Accurate et al.) for two hours, washed and suspended in a small volume for counting under the fluorescence microscope.

### 2.4 Immunohistochemistry

The hippocampi immunohistochemistry analysis was performed in 42-day-old NPP and NPE (n=5 for each group), and age-matched LPP and LPE (n=5 for each group) offspring subgroups were anesthetized as described above (75 mg.kg−1 body weight, i.p.) and xylazine (10 mg.kg−1 body weight, i.p.). They were monitoring the corneal reflex controlled anesthesia levels. Tissue perfusion was performed with a peristaltic perfusion pump, maintaining a mean pressure of 120 mmHg. Each animal was perfused for 15 minutes with 5% heparinized saline solution at room temperature and followed by 20 minutes with 4% paraformaldehyde solution. After perfusion, the brains were fixed by immersion in 4% paraformaldehyde for 2 hours. After fixation, the material was included in paraplast and, using a rotating microtome, 5 µm sections were collected on silanized slides. The brain histological sections adhered to silanized slides, which were deparaffinized and subsequently submitted to antigenic recovery in steam, in citrate buffer, pH 6.0 for 30 min. After the slides cooled down, they were washed in PBS (0.1M, pH 7.4) three times (5 minutes each). At the end of this process, the sections were incubated with primary antibodies SOX 2 Mouse (Sta. Cruz) with a dilution of 1:200 and Ki-67 rabbit (Abcam) with a dilution of 1:200, diluted in BSA 1%, overnight, under refrigeration. After washing with PBS (4 times at 5-minute intervals), the sections were exposed to the specific secondary antibody, conjugated with fluorochromes (Alexa 555 and 488) at room temperature for 2 hours. After successive washes with PBS, the slide was mounted with Vectashield containing DAPI. Image analysis was performed using a laser confocal microscope.

### 2.5. Data presentation and statistical analysis

All data were reported as mean ± standard deviation of mean (SD). Data obtained over time were analyzed using a one-way analysis of variance (ANOVA). When one-way ANOVA analysis indicated statistical differences between groups, Bonferroni’s contrast test performed post hoc comparisons between means. Comparisons between two groups were performed using a two-way repeated measure ANOVA, in which the first factor was protein content in a dam’s diet, and the second factor was time. When the interaction was significant, mean values were compared using Tukey’s post hoc analysis. Student’s t-tests were used to evaluate studies involving only two independent samples, within or between groups. Welch’s test was used to correct situations characterized by heteroscedasticity (different variances between groups). An animal’s survival lifetime was assessed using Mantel-Cox and Gehan-Breslow-Wilcoxon tests. GraphPad Prism 5.00 was used for data analysis (GraphPad et al., USA). The level of significance was set at p ≤ 0.05.

## 3. RESULTS

At birth, the male LP offspring (n=35) had lower body mass compared to NP progeny (NP: 6.48 ± 0.57 g, n=76 vs. LP: 6.09 ± 0.45 g, n=70; p<0.001; t=4,606, df=140,8) (Figure 1). The body masses difference between both groups was substantially enhanced being that the body mass gain in LP was lower compared to NP progeny, as shown by the results of body mass on the seventh day (NP: 17.4 ± 2.4 g, n=34 vs. LP: 10.8 ± 1.9g, n=32, p=0.0001, t=12.34, df=64), 14th (NP: 28.6 ± 4.5 g, n=67 vs. LP: 16.0 ± 2.0 g, n=62, p=0.0001, t=20.27, df=127), and 21st (NP: 49.6 ± 7.6 g, n=71 vs. LP: 22.3 ± 3.0g, n=63, p=0.0001, t=26.72, df=132) day of life (Figure 1). On the 42nd day of life, the difference between LP (n=40) and NP (n=40) offspring body mass was maintained (p=0.001). The NP progeny submitted to the enriched environment (NPE) showed an accentuated reduction in body mass compared to those rats kept in the standard environment (NPS: 182.0 ± 5.5g, n=5 vs. NPE: 158.2 ± 7.3g, n=5; t=5.823, df=8; p=0.0002). However, there was no significant difference between LPS and LPE offspring body mass (LPS: 118± 2.4 g vs. LPE: 113± 3.9 g, Figure 1).

**Figure 1.**
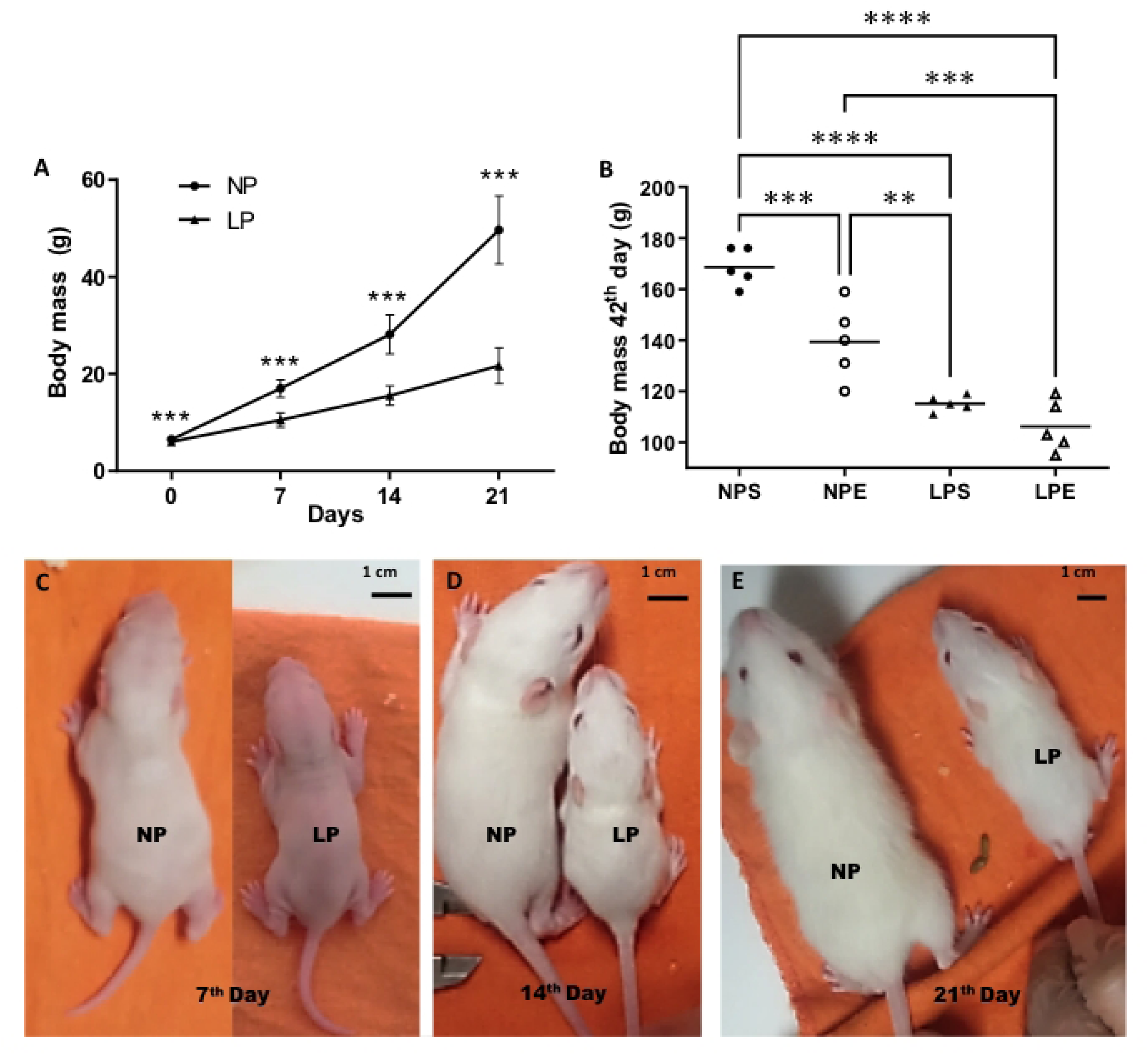
(A) Offspring body weight (g) at birth to 21 days of age (breastfeeding time), and (B) at 42-day-old NP (n=40), as compared to LP (n=40) offspring. The figure also includes representative photographs of the size difference between the NP and LP groups at 7, 14, and 21 days of extrauterine life. (Fig. 1 C-E, Bottom Panels). Our results, expressed as means ± SEM, were obtained over time and analyzed using repeated-measures ANOVA. Comparisons involving only two means within or between groups were performed using a Student’s t-test. Welch’s t-test was used to correct situations characterized by heteroscedasticity. When statistically significant differences were indicated between selected means, post hoc comparisons were performed using Bonferroni’s contrast test. The significance level was set at *P<0.05; *** P<0.0005.

On the 42nd day, we found no statistically significant differences in encephalic mass between the NP and LP groups when considering the ratio by body mass (NP: 0.568 ± 0.00972g; LP: 0.588 ± 0.00882; p=0.0632; t=1.595; df=20), or when comparing only the absolute values of the encephalic mass. Similarly, there was no difference in the ratio between hippocampal to brain mass (NP: 0.139 ± 0.0162g; LP: 0.169 ± 0.0136g; p=0.088; t=1462; df:9) for all groups. No significant difference was observed in the absolute hippocampal mass between the groups.

### Hippocampal total cell and neuron quantification and Immunohistochemistry

By the isotropic fractionation technique, a method that allows for the separation of different cell types in the hippocampus, by breaking down the tissue into isotropic fractions, the current study showed a difference in the hippocampal number of total cells comparing the NPS offspring (p=0.0170) with all of others experimental groups but not comparing the other groups with each other offspring groups (Figure 2A). On the other hand, the number of neurons was significantly reduced in the LP offspring (p=0.0001) compared to any of the NP progeny (p=0.0001); however, the enriched environment for three weeks (LPE) led to the reestablishment of neuron number comparing to age-matched NP progeny (NPS: 2.53 x 10^6^ ± 8.5.10^4^; NPE: 3.26 x 10^6^ ± 1.87 x 10^5^; LPS: 1.63 x 10^6^ ± 5.76 x 10^4^; LPE: 3.54 x 10^6^ ± 2.0 x 10^5^, n = 4 for each group, Figure 2B). Thus, it was observed that protein restriction during gestation and breastfeeding led to an increase in the ratio between neurons and other cell types Figure 2C, present in the hippocampus (non-neuron cells was relatively increased in LPS, with no statistical difference between groups, except to reduced NPS non-neuron cells, p=0.0415), and this was partially restored to that observed in the control animals after being in the enriched environment.

**Figure 2.**
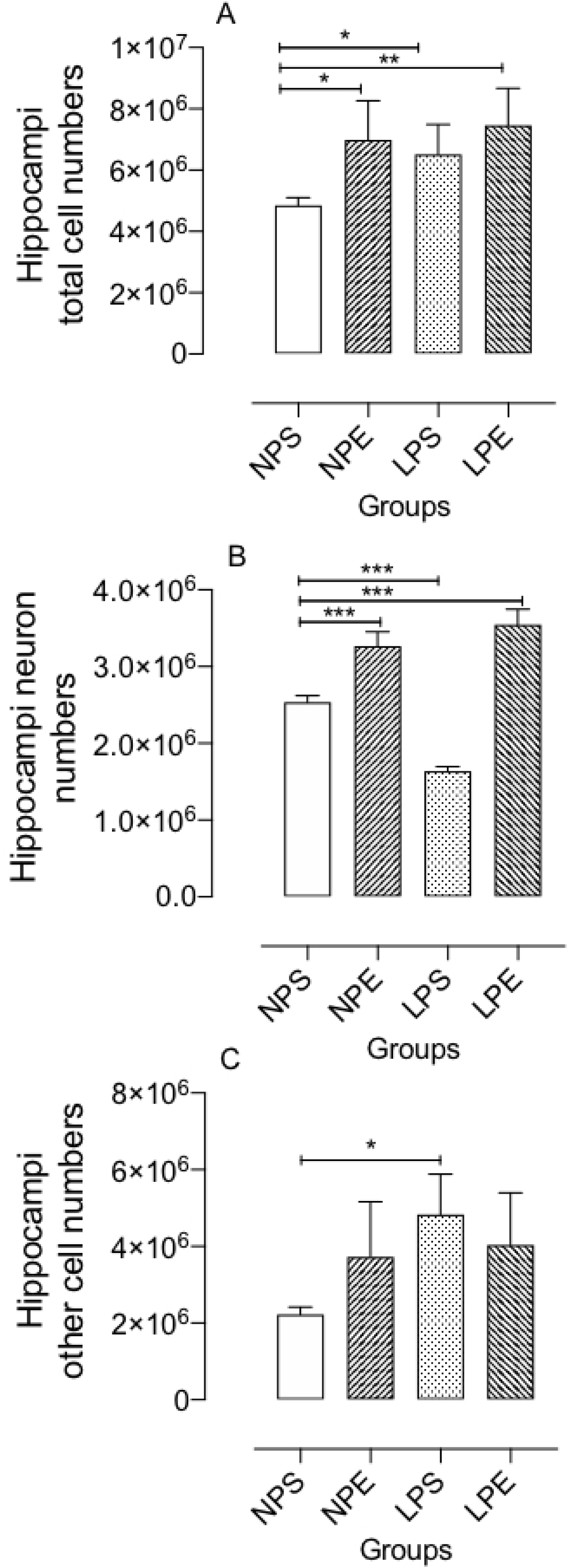
Representative graphics of the hippocampal effects of maternal protein restriction on 42-days-old LP (n = 5) compared to age-matched NP (n = 5) offspring on whole hippocampal cell (A), neurons (B), and non-neuronal cell (C) quantifications obtained by isotropic fractionation technique. Results are depicted as scatter dot-plot and are expressed as means ± SEM; comparisons involving only two means within or between groups were performed using a Student’s t-test. The level of significance was set at *p < 0.05.

Also, the immunohistochemistry technique of hippocampus regions evaluated the number of mitoses and stem cells in the subgranular and granular areas of the dentate gyrus (n=5 for each group). An expressive and significant reduction of cells in mitosis, stem cells, and stem cell mitosis was observed in the LP offspring in the granular layer (GL) and subgranular zone (SGZ) of the DG (Figure 3). These results demonstrated that the exposure of animals to the enriched environment promoted the retrieval of the cell proliferation process, particularly of stem cells (Figure 4). These observations were also evidenced in the subventricular zone near the lateral ventricles. They presented statistical significance by quantification (Figures 4 and 5).

**Figure 3.**
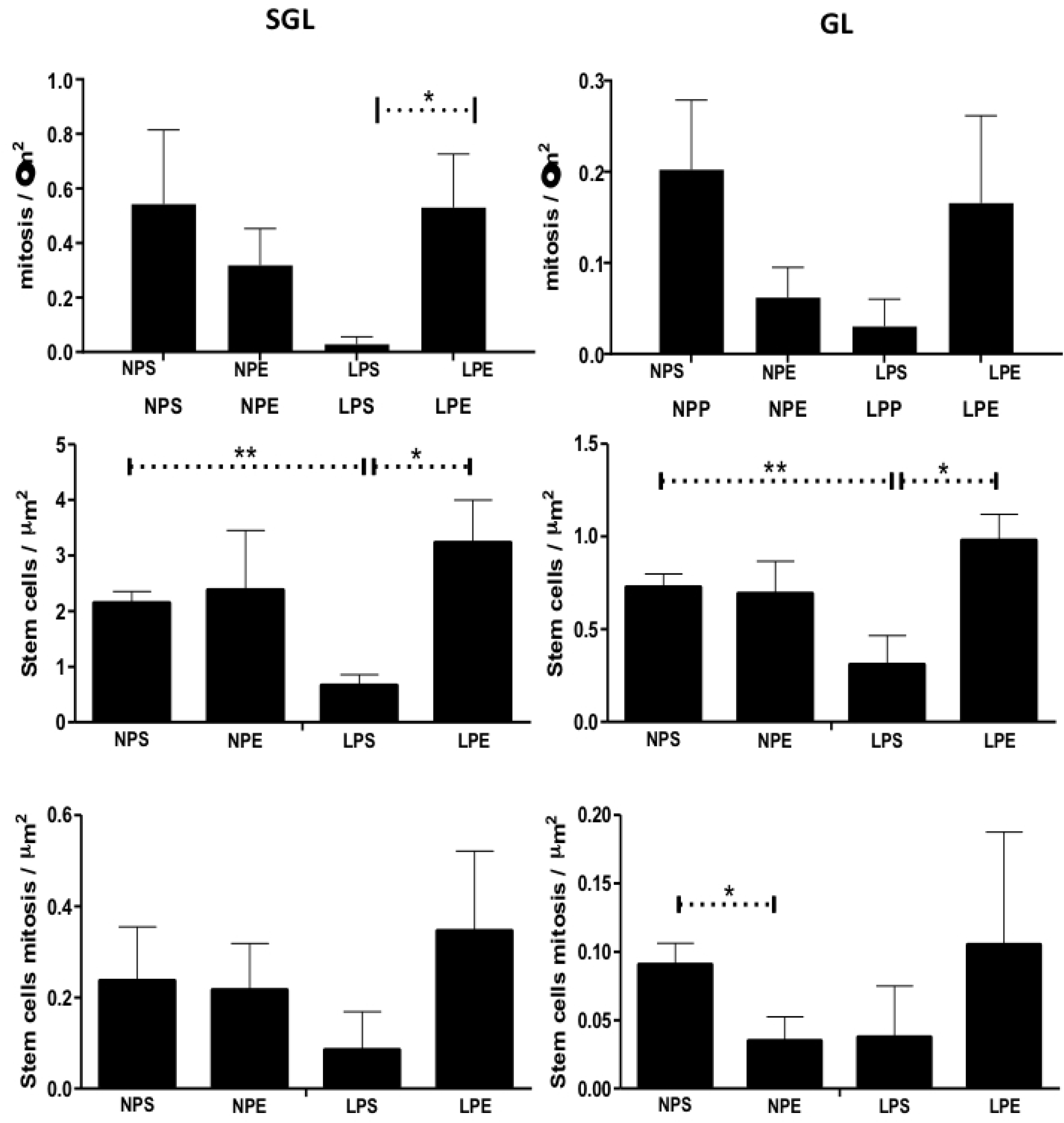
Representative graphics of the number of mitoses, stem cells, and stem cell mitoses quantified in the subgranular (SGL) and granular (GL) layers of the hippocampal dentate gyrus (GD) in the offspring. On the left of the panels are presented the groups of offspring studied whose mothers were subjected (LPS) or not (NPS) to protein restriction during pregnancy and breastfeeding and exposed to an enriched environment after breastfeeding (LPE and NPE). Results are expressed as means ± SEM; the significance level was set at *p < 0.05 (one-way ANOVA analysis indicated statistical differences between groups, Bonferroni’s contrast test performed post hoc comparisons between means); n=4 for each group.

**Figure 4.**
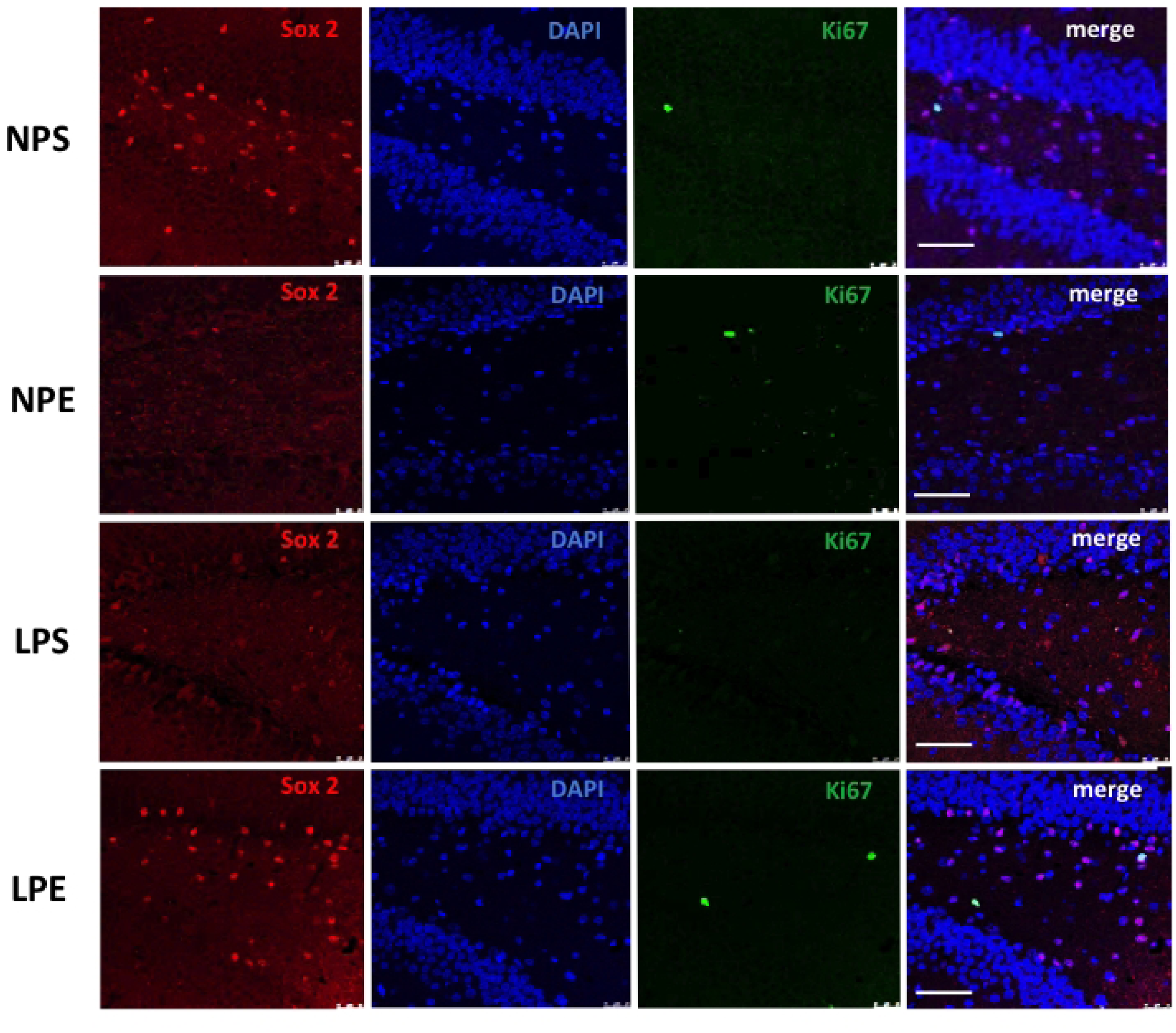
Representative images of hippocampus regions were obtained using immunohistochemistry, evaluating the number of mitoses and stem cells in the subgranular and granular areas of the dentate gyrus (n=5 for each group). On the left of the panels are presented the groups of offspring studied whose mothers were subjected (LPS) or not (NPS) to protein restriction during pregnancy and breastfeeding and exposed to an enriched environment after breastfeeding (LPE and NPE).

**Figure 5.**
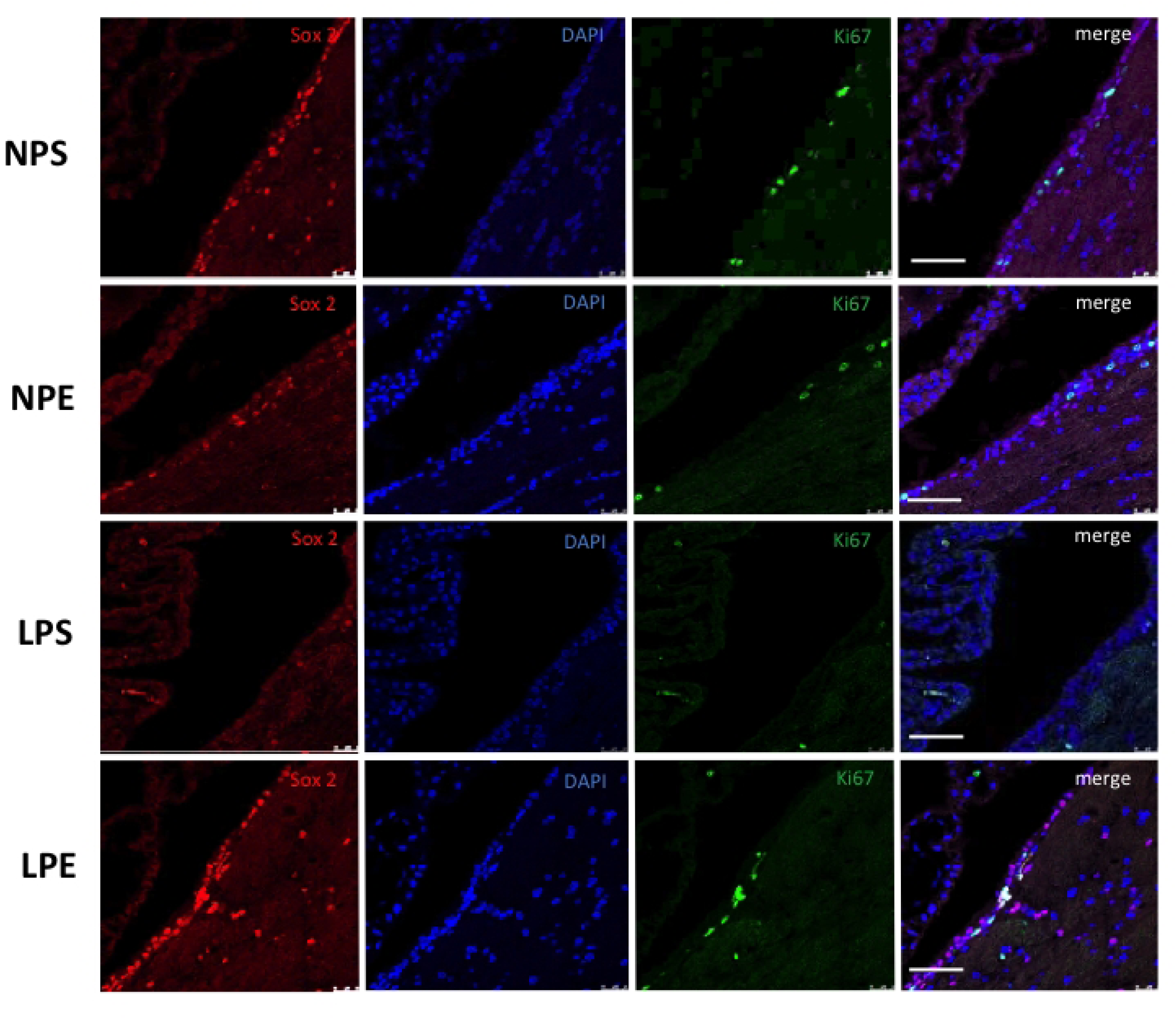
Representative images obtained by the immunohistochemistry technique of subventricular regions near the lateral ventricles, evaluating the number of mitoses and stem cells. (N=5 for each group). On the left of the panels are presented the groups of offspring studied whose mothers were subjected (LPS) or not (NPS) to protein restriction during pregnancy and breastfeeding and exposed to an enriched environment after breastfeeding (LPE and NPE).

### Behavioral Tests

#### Activity Monitoring Test (Open field tests)

The distance covered by the animals was considered the sum of the different displacements of positions monitored by the equipment. The key finding of this test is that the enriched environment significantly increased motor activity in NP animals. Although a slight increase in LPE was observed in offspring, the difference was insignificant compared to LPS (Figure 6, Table 2). The average speed of the animals was calculated as the distance traveled from one point to another, divided by the time taken by the animal to cover this distance. Another significant finding is that there was a considerable increase in NPE animals compared to NPS, and there was no substantial difference between LPS and LPE (Figure 6, Table 2). As for motor activity, jumping, rearing, rotational and ambulatory movements, and times standing in an orthostatic exploratory position, the observed behaviors were similar for all groups studied. However, being greater in NPE offspring compared to control and LPE progeny. This way, an increased result in these behavior parameters is observed only in NP submitted to the enriched environment (Table 2). In isolation, the number of times animals assumed orthostatic positions showed a more minor but insignificant comparison between LPS and NPS (Table 2). The stereotyped movements of these animals were considered when showing a scratching attitude, repetitive masticatory, or intensive tail movements. The LP progeny did not show a change in stereotyped movements compared to NP offspring, and the enriched environment did not promote any change in this parameter (Table 2). The time spent by animals remaining in the center or on the arena’s edges was distinct. The LP offspring stayed longer at the edges and avoided the arena center (Table 2) relative to NP rats. The enriched environment unchanged this behavior in both NP and LP offspring (Table 2).

**Figure 6.**
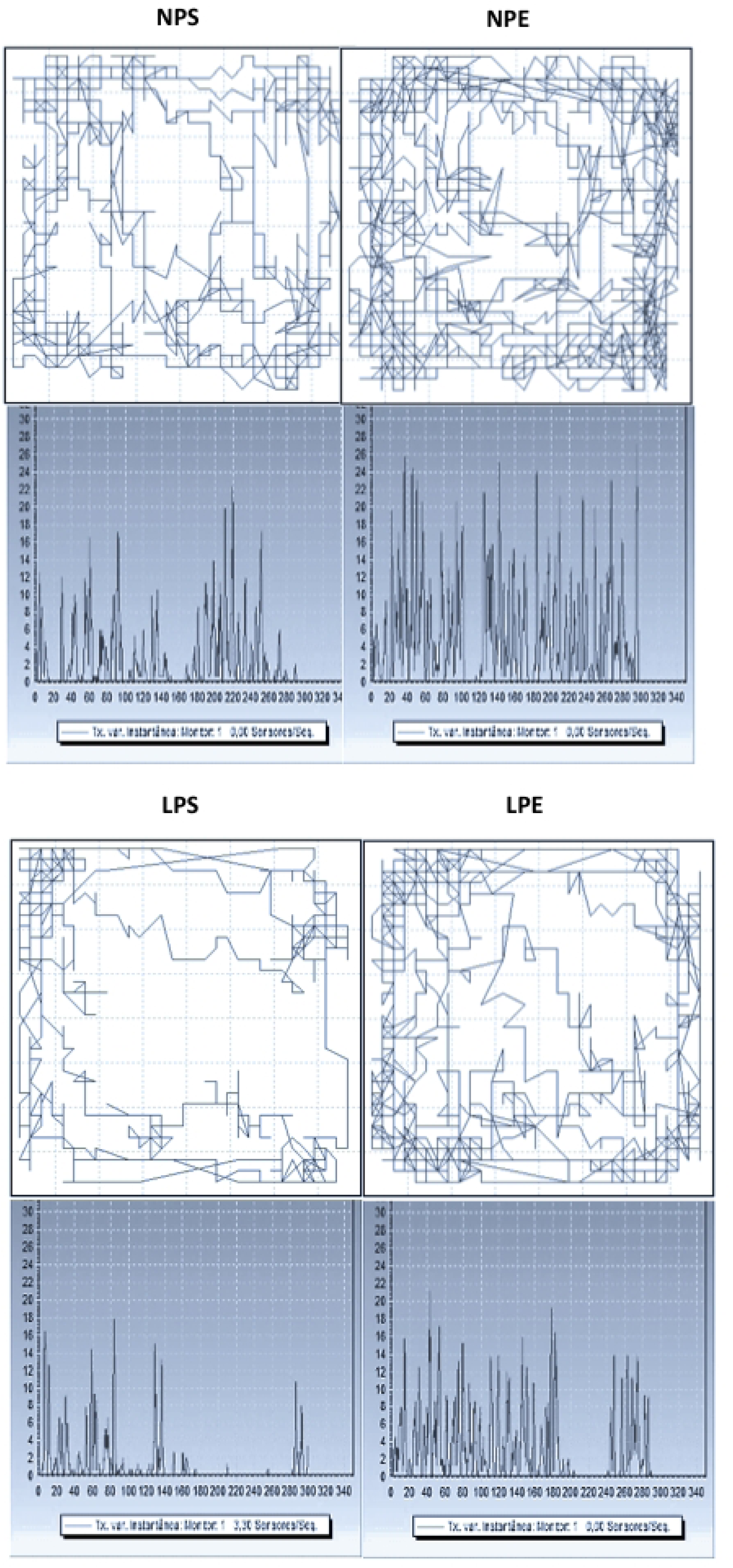
Representative results were obtained by the activity monitor in the open field test and the level of activities. In the upper images, we have represented examples of the displacements of NP progeny keeping in a standard environment (NPS) and the other in an enriched environment (NPE), and the bellow images are representative of displacements of LP offspring in a standard (LPS) or enriched environment (LPE). In the lower images, we have examples of the rate of variation of activities per second for these animals.

**Table 2.**
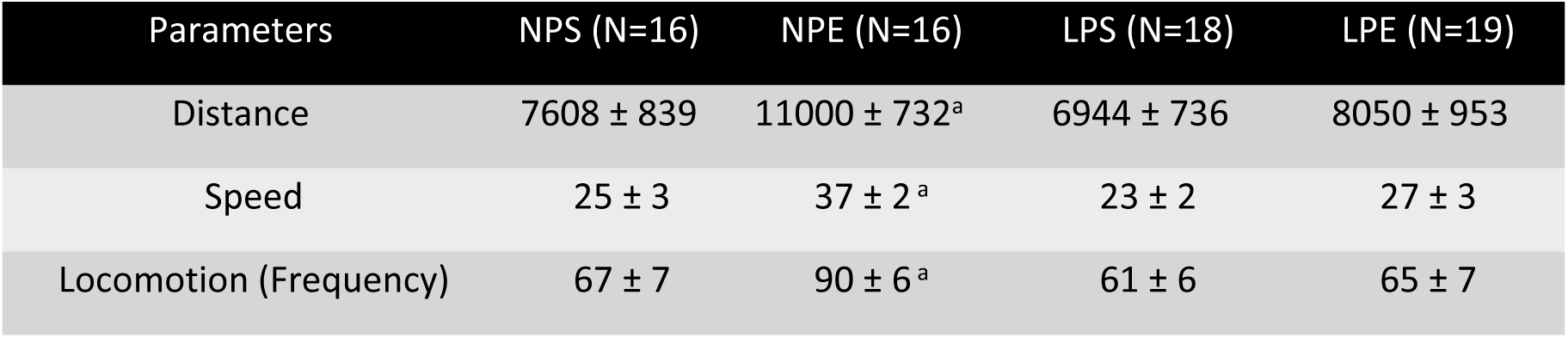
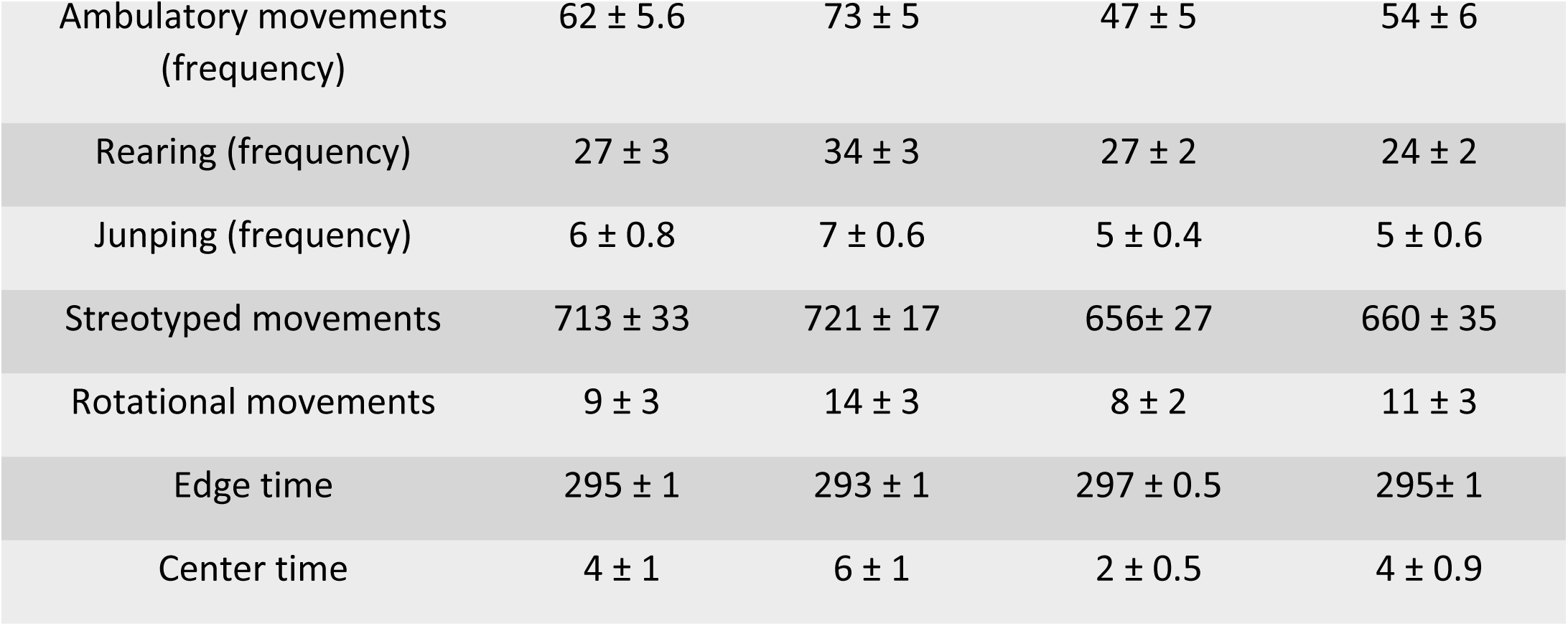
Open field results from 16-week-old regular protein diet (NP) (n =13) v. age-matched low protein content (LP) (n =13) offspring. The results are the mean frequency of rearing, dislocation, freeze, self-grooming, and crossing throughout the open field. (Student t-test; a: NPS vs. NPE; b: LPS vs. LPE; c: NPS vs. LPS; d: NPE vs. LPE).

The number of fecal boluses in NPS or LPS did not change during the five minutes of observation; however, the enriched environment significantly decreased the number of fecal boluses, indicating a reduction in fear-reflecting behavior in both groups (Figure 7).

**Figure 7.**
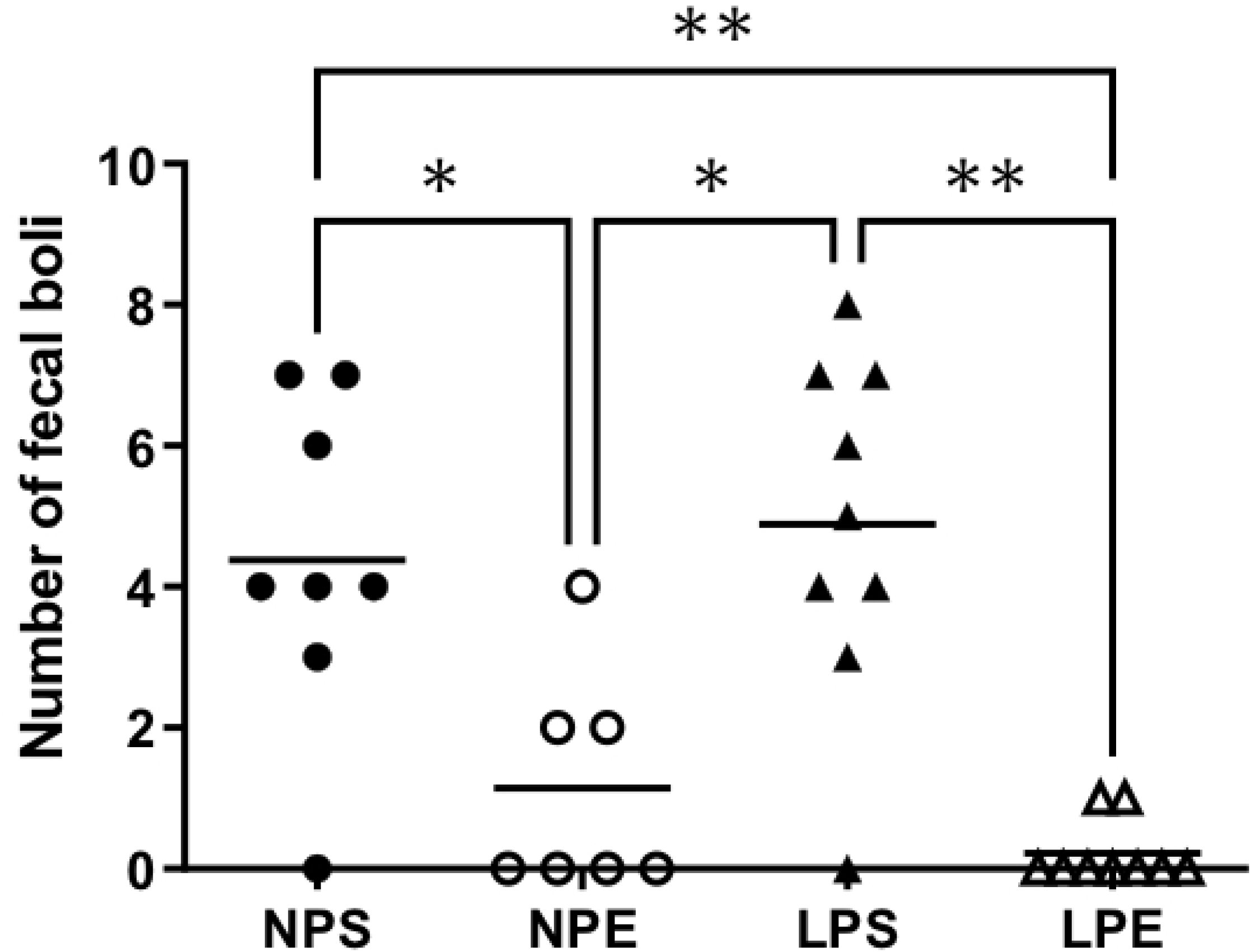
The mean percentage of open arm time (A) and open arm entries (B) for rats in the elevated plus maze (EPM) is depicted. The exploration in the elevated plus maze was meticulously measured. The enriched environment significantly increased the time spent and entries in the open arms of NP and LP animals. Notably, only LPE showed a significant rise. This finding underscores the impact of environmental enrichment on animal behavior. * Indicates p<0.05 and ** p<0.0005 (one-way ANOVA analysis indicated statistical differences between groups, Bonferroni’s contrast test performed post hoc comparisons between means); n=17 offspring for each group.

#### Elevated plus maze test

The animals in the NPS group remained significantly perceptual longer time in the center point of the apparatus (NPS: 23 ±1.2%; NPE: 16.8 ± 1%; LPS: 15 ± 1.8%; LPE: 14.3 ± 1%, Figure 8) compared to other groups. However, no difference was observed between non-stimulated NP and LP regarding permanence or entry into the open arms. After the enriched environment exposure, an increased permanence or entry into the open arms in both groups (NPS: 13.8 ± 9%; NPE: 21.7 ± 11%; LPS: 14.8 ± 10%; LPE: 26.3 ± 7.2%, Figure 8) was observed, indicating a decrease in anxiety-simile behavior in this situation.

**Figure 8.**
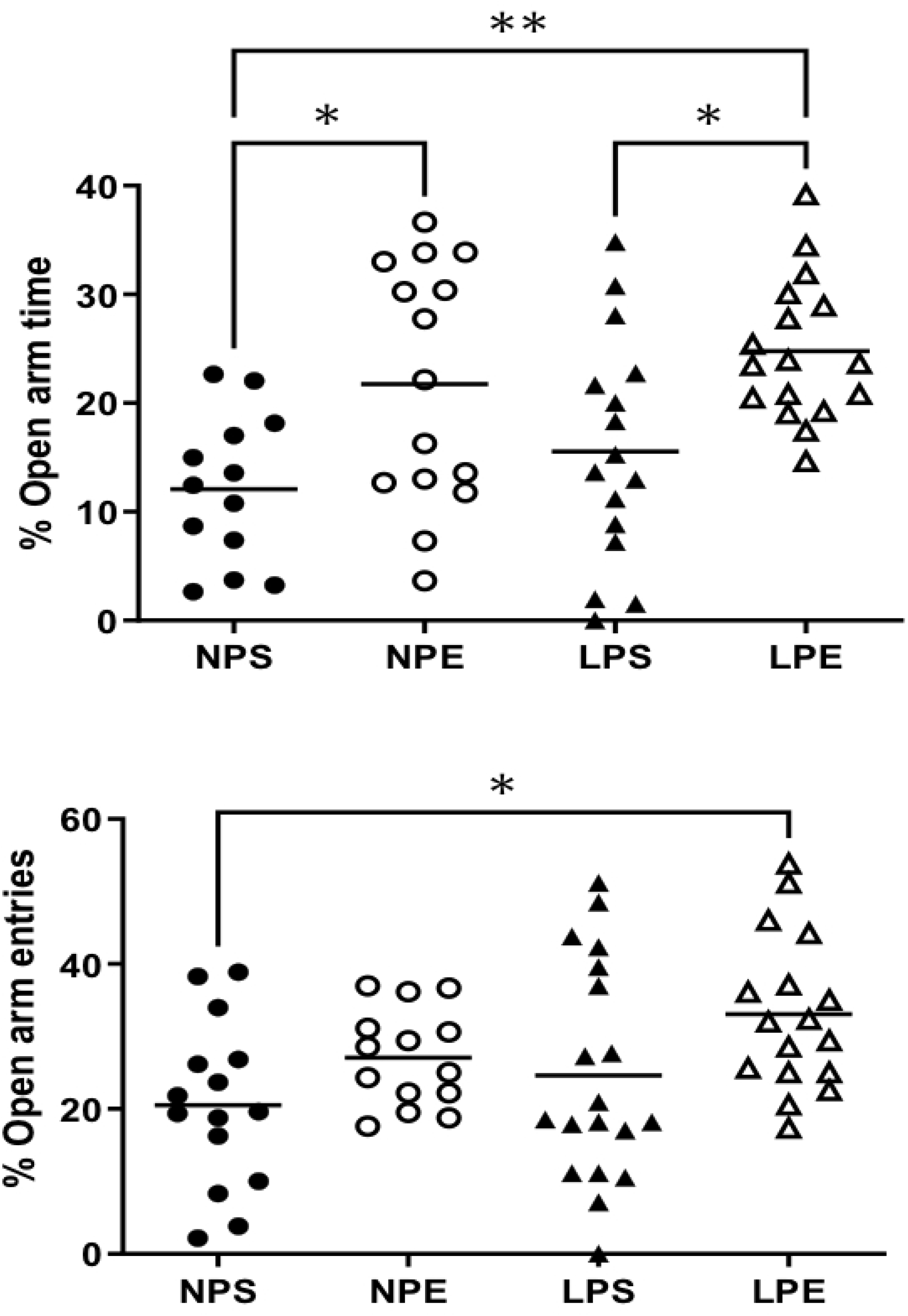
The figure presents the dot-plot of fecal cakes as means ± SEM to NP and LP animals subjected to standard (NPS, LPS) or enriched environment (NPE, LPE). The LPS and LPE offspring notably exhibited a significantly smaller number of fecal pellets, indicating a potential improvement in their health. This stark contrast between the standard and enriched environments underscores the role of environmental factors in animal well-being. *P< 0.05; ** P<0.005; ***P<0.0005 (one-way ANOVA analysis indicated statistical differences between groups, Bonferroni’s contrast test performed post hoc comparisons between means); n=17 offspring for each group.

#### Novel object discrimination test

The enriched environment significantly modified the short-term discrimination ratio, increasing the ability of NP and LP animals to discriminate between novel and familiar objects quickly (Figure 9). However, we did not observe a statistical difference between the groups in the long term.

**Figure 9.**
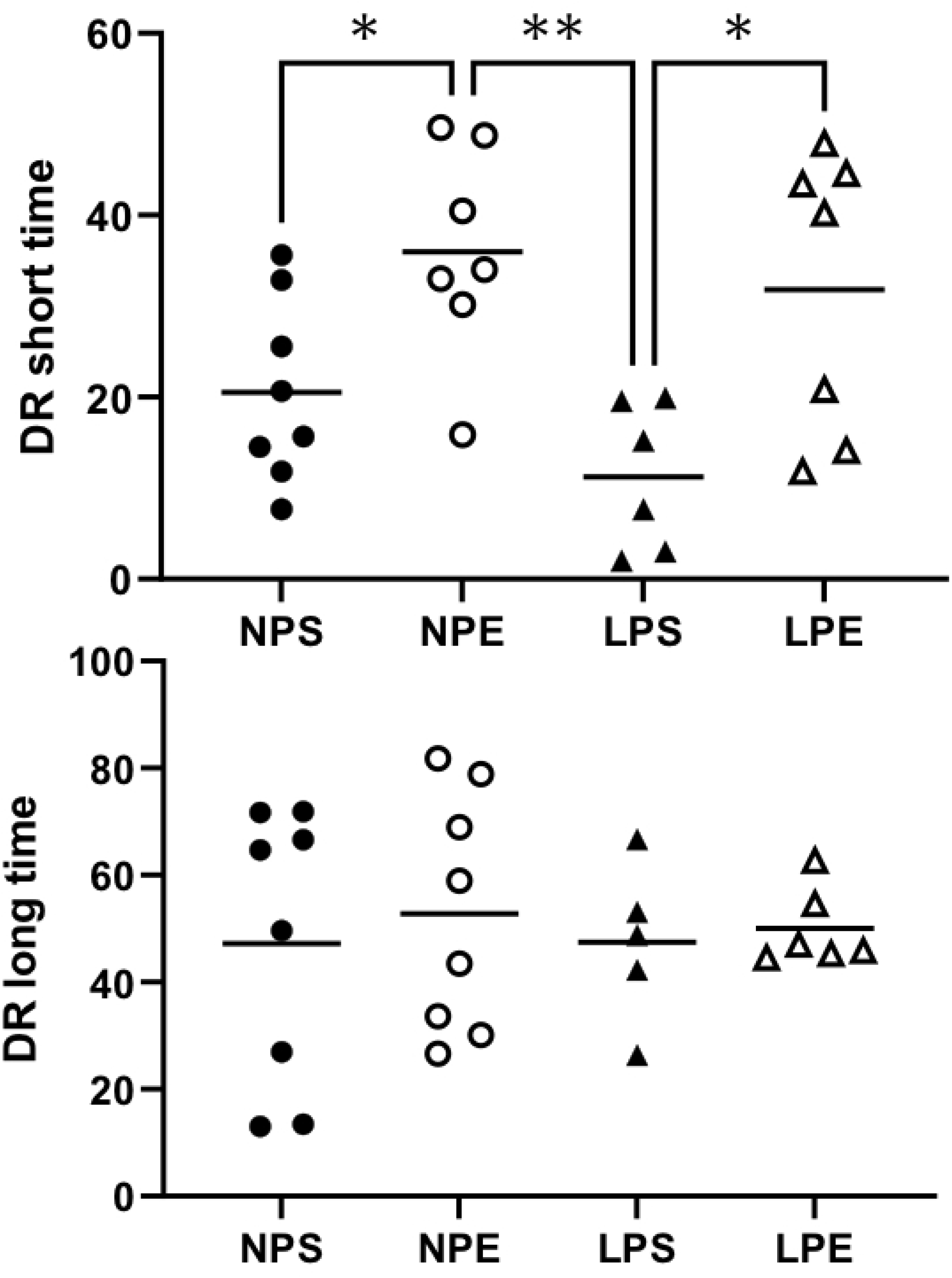
The graph represents the new and familiar object discrimination ratio (DR) on short (A) and long (B) time presented by the four groups evaluated. *P< 0.05; ** P<0.005 (one-way ANOVA analysis indicated statistical differences between groups, Bonferroni’s contrast test performed post hoc comparisons between means); n=5-8 offspring for each group.

#### Morris water maze

Our study found a significant difference in the learning process between the NP and LP groups in trials 2 and 3 (F = 71.89, p = 0.0001). This difference was particularly pronounced across days, suggesting an enhanced learning process in LP offspring (p = 0.001; Figure 10A). For the reference memory, repeated-measures ANOVA for escape latency showed reduced escape latency in the LPE group on days 1 and 2. Again, both groups learned the task across the days (NP: 31.53 ± 8.48 vs. LP: 24.27 ± 7.71; p=0.0033, t=2.868, df=39; Figure 10C). On the eighth day, the escape platform was removed, and the LPE animals demonstrated superior memory retention by remaining longer in the quadrant where the platform was on the previous day, compared to the NPS and NPE groups (Figure 10C).

**Figure 10.**
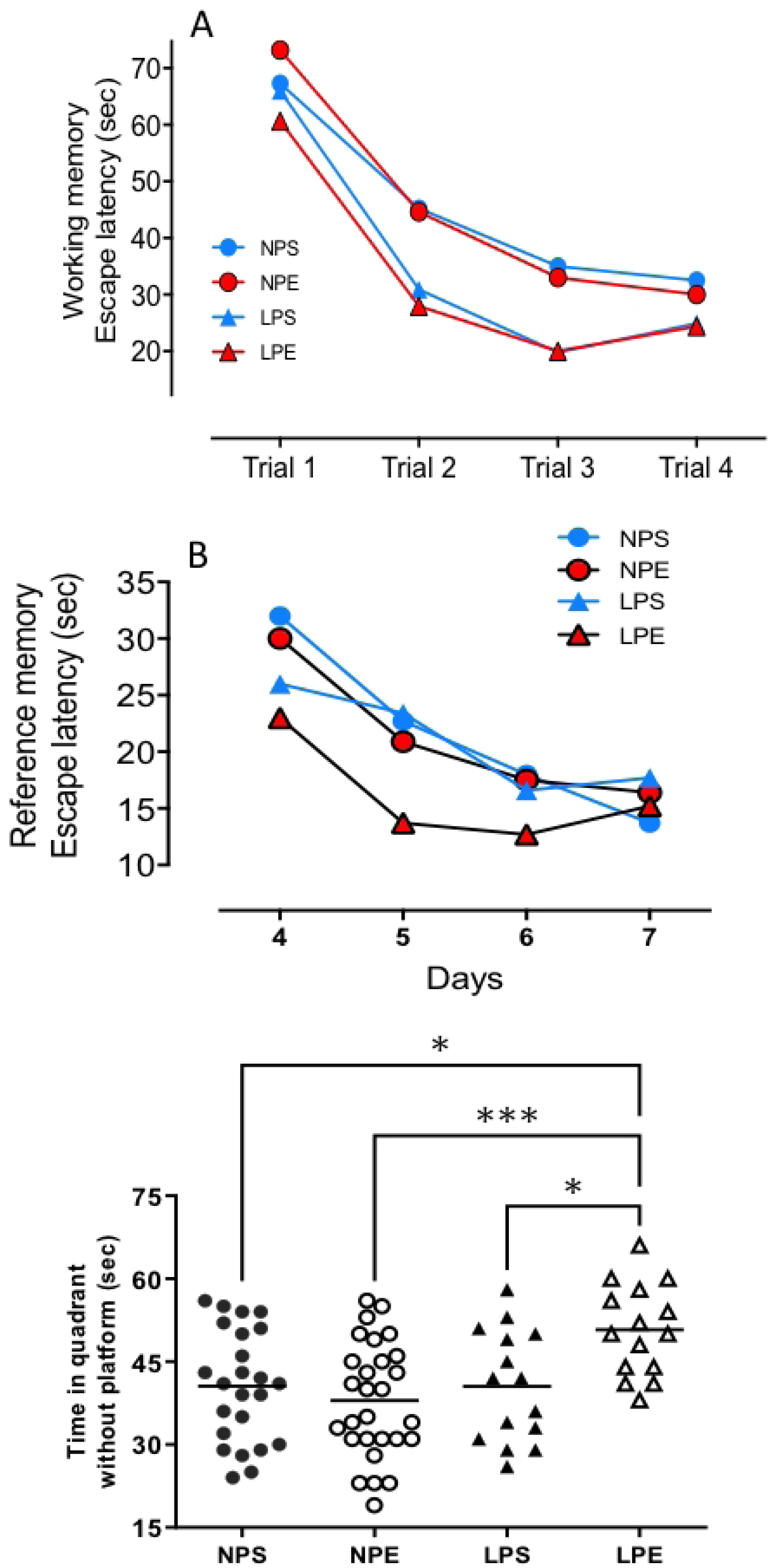
The Morris water maze training performance is illustrated. (A) In the working memory test, the LP animals demonstrated a faster learning rate than the NP, indicating a potential cognitive advantage of the LP diet. (B) The reference memory test revealed that LPE animals learn faster than others, highlighting the enriched environment’s cognitive benefits. (C) As memory retention results, on the eighth day, the LPE animals remained significantly longer in the quadrant where the platform was on the previous day than NPS and NPE, suggesting a superior memory retention ability.*indicates p < 0.05 and**p < 0.01.

## 4. DISCUSSION

The topic of the current study, undernutrition in perinatal life, is essential as it considerably modifies the development and maturation of tissues and organs in developed and underdeveloped countries [43,44]. Malnutrition may cause structural and functional changes in organs and systems, including the CNS, partly due to increased fetal exposure to maternal glucocorticoids, which alter the synapses and neuron dendritic branching and number [45,46]. The hippocampus development in mammals has received increasing attention due to its significant neuroplasticity and structural and functional abnormalities resulting in the offspring’s emotional (anxiety/fear), learning, and memory disorders [26,47,48].

Our study consistently observed a reduction in birthweight in the offspring of protein-restricted mothers, a trend that persisted and intensified post-natally compared to control litters. Interestingly, despite these changes, the brain mass in the LP offspring remained unchanged. This suggests that the offspring’s peripheral organs adapt by decreasing energy metabolism to maintain a stable brain energy supply, which aligns with the known selfish brain theory in these offspring. Furthermore, the study revealed a significant change in hippocampal cell composition in the LP offspring, indicating a potential adaptation to the protein-restricted environment.

The current study aimed to evaluate the effects of pregnancy and breastfeeding protein intake restriction on the structure and function of the hippocampus and the relationship between hippocampus cell pattern and potential behavioral changes, including after exposure of male offspring to the enriched environment (EE). Here, the findings confirm prior studies showing that malnutrition, when imposed during the period of CNS development, both in the gestational and lactation periods, affects mainly the body mass, regardless of the environmental stimulation to which the animal is subsequently subjected [1,4,49–51], have significant implications. Surprisingly, after breastfeeding, the EE promoted an additional reduction in body mass in offspring with average protein intake (NP) and offspring with low protein intake (LP). In addition, the pregnancy and lactation protein restriction also caused the decrease in hippocampus neurons by non-neuron cell ratio, and the regular proportion has been restored compared to age-matched NP progeny after exposure of the LP offspring to the EE for three weeks.

Our study uncovers a novel aspect of the hippocampus constitution, where the functional significance of this cellular pattern change in LP offspring remains a mystery. This is the first instance where changes in the hippocampi cell pattern, followed by cytological restoration through exposure to EE, have been demonstrated in the maternal protein-restricted offspring. While these results are unprecedented in LP rats, a previous study has shown increased brain mass, thickening cortical layers, and biochemical and cellular constitution after EE exposure [52,53]. Furthermore, supporting our present data, brain plasticity studies have demonstrated increased neurogenesis [53], gliogenesis, and dendritic arborization [54] in mice after EE exposure. Authors have reported an increased synaptic density in environmentally stimulated animals [52, 55] associated with increased cortical expression of genes, such as the Arc gene, involved in the molecular process of cellular plasticity [56]. These EE-induced changes were associated, at least in part, with the decreased cell death by apoptosis in the hippocampal region, the protective effect on aging, and the increase in neurotrophic factors [37,54,57,58].

The present study revealed an imparity between behavioral test response and changes in the hippocampal neuronal number, with increased non-neuronal cells, as a consequence of fetal programming. Therefore, this study did not demonstrate a difference in the whole hippocampal cell number in the LP offspring compared to the age-matched NP group by isotropic fractionation technique. However, the number of stem cells and neurons was reduced in LP progeny. This data suggests a concomitant increase in non-neuron hippocampal cells, which, in turn, maintain an even hippocampus mass relative to NP progeny.

Immunohistochemistry confirmed these findings and demonstrated that the number of stem cell mitoses in the SGZ and GL of the hippocampal DG in the offspring of NP and LP were highly distinct. Thus, a reduced number of progenitor cells, neurons, and total cells in mitosis were observed in the GL and SGZ of the DG in the LP offspring. However, exposing male LP offspring to the enriched environment significantly increased hippocampal cell proliferation, thereby recovering the neuronal stem cell number. This methodological approach allowed us to observe and quantify the changes in the hippocampal cell composition, providing valuable insights into the effects of maternal protein restriction and the potential of environmental enrichment to reverse these effects.

The present data, in line with prior studies, confirm that gestational protein restriction poses a threat to brain development, reducing dendritic ramifications, the branching out of dendrites that allow neurons to communicate with each other, the number of synaptic terminals, the points of contact between neurons where information is transmitted, altering the neurochemistry, and myelination throughout the CNS [6–8,41,42]. Additionally, studies have shown that maternal chronic stress and corticosteroid treatment induce dendrite atrophy, a condition where dendrites shrink or degenerate, of pyramidal neurons with decreased length and the branching points of the apical neurons in hippocampus CA1 and CA3 [59–67]. These findings underscore the urgency of understanding the effects of maternal protein restriction on offspring’s hippocampus development and behavior and the potential for environmental enrichment to reverse these effects partially.

The high glucosteroid receptor (GR) mRNA’s expression in the hypothalamic paraventricular nuclei (PVN) is widely recognized; otherwise, the mineralocorticoid receptor (MR) gene expression is present exclusively in the limbic system, hippocampus, amygdala, and DG [68]. In rodents, maternal adverse events during gestation alter the hippocampus-HPA axis activity, a complex interaction between the hippocampus and the hypothalamic-pituitary-adrenal (HPA) axis that regulates stress response, accompanied by GR down-regulation and an exacerbated response to stressful stimuli. It has been demonstrated that MR mediates hippocampal neurogenesis, the process of generating new neurons in the hippocampus. GR suppresses neuronal differentiation, the process by which a neuron becomes specialized for its function [60,69]. Studies in our Lab comprised an increased serum corticosterone level beyond seven days of age in LP offspring compared to NP progeny [7,8]; so, we may suggest that increased serum steroid level is a crucial factor involved in the structural hippocampus remodeling, characterized by dendritic atrophy of CA3 as well a reduction in the neuron number.

Here, seeking to associate hippocampal morphological findings with behavioral evaluation, the NP and LP offspring were submitted to activity monitoring, elevated plus maze, and the new object discrimination tests. Despite the changes in hippocampal cellular pattern identified in the LP offspring with an essential reduction in neuron numbers, no difference in recognition memory was found compared to the NP progeny. However, exposure of animals to EE caused enhanced short-term recognition memory capacity in both offspring groups, which was more pronounced in the LP and associated with an increased number and differentiation of the DG neurons. Here, it is essential to emphasize the role of motor activity and exploration capacity in the acquisition phase of recognition memory tests. Thus, reduced motor activity, or the little exploration in LP offspring, may compromise the recognition memory capacity. The monitoring activity showed no differences in locomotor activity caused by maternal protein restriction; however, the EE increased the NP offspring’s average speed and motor activity without any change in the LP offspring. Additionally, the EE causes unchanged stereotyped and rotational movements of the NP and LP animals. These findings suggest that the differences in the cytological hippocampal patterns observed in LP progeny do not substantially modify the motor response in this experimental model.

As previously stated, the hippocampus is critical in memory/cognition and emotional control [70]. Our study, however, brings a fresh perspective. We found that environmental enrichment (EE) significantly impacted the discrimination capacity test, enhancing the ability of both NP and LP offspring to distinguish new from familiar objects. Interestingly, this effect was more pronounced in maternal protein-restricted offspring in the short term, with no discernible difference in the long term between the groups.

Berlyne, in the fifties, described rats as spending more time exploring new than familiar objects [71]. In this way, Ennaceur and Delacour (1988, 1995)[72,73] and Forwood et al. (2007)[74] associated the animals’ preference for object novelty in different sites with previously found recognition memory capacity. Also, studies have demonstrated an altered capacity of animals concerning recognition memory, indicating a fundamental role of the hippocampus in storing information during the memory process [75–80]. In coincidence to hippocampus cytological changes from the current study, Good et al. (2007)[78] and Caceres et al. (2010)[79] caused neuron death, respectively, by intracerebroventricular ibotenic acid microinjection and radiation in rats promoting about 70% hippocampal neurons loss with significant compromised recognition memory. These animals had a lower object recognition index compared to the control group.

Our study, as confirmed by the present findings, demonstrates the significant impact of EE on behavioral (cognitive, sensory, motor, and affective tests) and memory tests. This improvement is a direct result of the increased number of neuron synapses and postsynaptic density in the hippocampus and parietal cortex [52,81–83]. The habituation assessed through the open field test [84–85], startle reflex [86], and elevated plus maze and eight-armed radial maze [87] further supports these findings. As reported by the authors, the benefits of EE can last for several months and are directly proportional to the exposure time. However, it’s important to note that the effects of the enriched environment on the brain are diverse, as numerous models of EE vary in terms of exposure time and the age at which exposure begins.

The EPM is the most used animal test for studying fear/anxiety by several research groups worldwide. In the present study, although we did not observe a significant difference between NP and LP results regarding permanence and entry into the open arms, the EE exposure significantly increased the permanence and numbers of entries in open arms in both NP and LP progeny, indicating a reduction in fear/anxiety-simile Behavior by EE, particularly in LP offspring. It is assumed that the open arms of the maze combine two components that are naturally aversive to animals: a new environment and an open space without protective walls. Nonetheless, in the current study, the LP offspring were unable to show changes in the entry numbers or time spent in the EPM open arms compared to the NP offspring despite extensive hippocampal cytological alteration. On the other hand, the LP offspring submitted to EE remained in closed arms for less time than in open arms compared to non-stimulated animals. These findings suggest an EE protective effect against possible behavioral changes induced by protein malnutrition. This protective effect of EE was associated with hippocampal cell pattern restoration, suggesting environmental stimulation might play a role in brain plasticity [88,89] and, concomitantly, in behavior response tests [56,89]. Several studies have reported improved motor and cognitive recovery of animals with ischemic brain injuries and underwent EE associated with increased astrocytes, oligodendrocytes, and neural stem cells [91–93]. Also, in experimental models of epilepsy, EE promoted increased resistance to seizures and decreased cognitive deficits, accompanied by changes in the cellularity pattern [37,84,94]. In the current study, there was no change in the number of fecal boluses in NP or LP during the five minutes of observation; however, the enriched environment significantly decreased the number of fecal boluses, indicating a reduction in fear-reflecting behavior in both groups.

Building on previous work in our laboratory, our study uses the Morris water maze test in the same experimental model. However, this time, incorporating gestational protein restriction has further demonstrated spatial reference memory’s unique and crucial role in the hippocampus. This finding underscores the importance of our study in contributing to the understanding of brain plasticity and behavior response tests [6].

Both the hippocampus and the prefrontal cortex are involved in spatial memory processing [95]. Surprisingly, despite the cytological changes, the present study did not find significant differences between the NP and LP offspring in any analyzed behavior tests. In a similar model of gestational protein restriction, we also did not observe modifications in spatial memory, learning, or working memory [15,96]. Previously, our lab’s study had detected a dissociation between function (learning and memory) and hippocampal structure, which was evaluated by the three-dimensional Golgi-Cox staining analysis in the dorsal hippocampus. It was demonstrated that gestational protein restriction leads to a decrease of about 30% in the whole length of basal dendrites and the number of intersections of apical dendrites from pyramidal neurons of CA3. In this study, the dendritic architecture of the DG and CA1 neurons remained unchanged [6].

In the present study, we observed a significant worsening in LP’s working memory compared to NP, and exposure to EE did not have any effect. In reference memory, despite changes in hippocampal cellularity, we did not observe changes in this parameter related to the LP diet during pregnancy and breastfeeding. However, EE improved reference memory only in LP animals.

The reference memory and retention test shows a difference in the escape latency between the groups, suggesting that LPE performed significantly better than controls. The probe trial results on the last day (Day 8) show that LPE rats spend considerably more time in the quadrant where the hidden platform was previously placed compared to the control. These data indicate that EE substantially improves memory in LP animals.

In conclusion, the current study findings have potential implications worth exploring. We demonstrated that maternal protein restriction during neural development causes crucial morphological changes in the hippocampus, making the LP offspring vulnerable to neural disorders in adulthood. This sustains the ‘selfish brain’ theory, as the paradigm that postulates the brain maintains its mass ‘selfishly.’ However, the hippocampus cellularity pattern was profoundly altered, significantly reducing the number of neurons after the breastfeeding period. We also observed reciprocal data, indicating a direct relationship between brain masses, changes in the hippocampus cell pattern, and decreased body mass in the LP progeny. We also demonstrated that neuronal composition and structure profoundly modified by dietary restriction are surprisingly restored from primordial cells by exposure to an enriched environment, which included increased social interaction, physical activity, and cognitive stimulation. Importantly, we observed a significant reduction in neurons after gestational and breastfeeding periods and, for the first time, a substantial reduction in fear-reflecting behavior, which an enriched environment exposure may revert. An enriched environment can partially mitigate these effects, highlighting the importance of early-life nutrition and environmental stimulation in shaping brain development and behavior. The enriched environment also significantly modified the discrimination ratio, increasing the ability of both progenies to discriminate between novel and familiar objects in a short time associated with reverse abnormal hippocampus cell patterns. EE did not reverse the harmful effects of the diet on working memory but significantly improved reference memory in LP. Here, we demonstrated that environmental enrichment might attenuate some of the deficits caused by protein restriction during gestation and breastfeeding.

## Acknowledgments and Financial support

This work was supported by Fundação de Amparo à Pesquisa do Estado de São Paulo (2013/12486-5), Coordenação de Aperfeiçoamento de Pessoal de Nível Superior (CAPES) and Conselho Nacional de Desenvolvimento Científico e Tecnológico (CNPq, 465699/2014-6).

Availability of data and material in: http://repositorio.unicamp.br/jspui/handle/REPOSIP/312739?mode=full http://repositorio.unicamp.br/jspui/bitstream/REPOSIP/312739/1/Grigoletti_GabrielBoerLima_M.pdf

## 6. ABBREVIATIONS AND ACRONYMS

11β-HSD2: type-2 11β-hidroxiesteroide dehydrogenase
ACTH: Adrenocorticotrophic hormone.
ANOVA two-way: Two-way analysis of variance
BDNF: Brain-derived neurotrophic factor
CEMIB: Center of Bioterism of the State University of Campinas
CEUA/Unicamp: State Campinas University Institutional Ethics Committee
CNS: Central nervous system
COBEA: Brazilian College of the Animal Experimentation
CORT: Corticosterone
CRF: Corticotrophin-releasing factor
CRF1: type-1 CRF receptor
CRH: Corticotropin-releasing hormone
CRHR: CRH receptor (CRHR1 and CRHR2)
DCX: Doublecortin
DG: Dentate Gyrus
DH: Dorsal Hippocampus
DNA: Desoxyribonucleic acid
EE: Enriched environment
EP: Epinephrine
EPM: Elevated plus maze
G.C.: Glucocorticoid
GABA: Gamma-aminobutyric acid
GR: Glucocorticoid receptor
GZ: Granular zone
Hippocampus-HPA: Hippocampus-hypothalamic-pituitary-adrenal axis
HPA axis: Hypothalamic-pituitary-adrenal axis
LP: Low protein diet content or LP progeny
LPE: Low protein diet content or LP progeny submitted to the enriched environment
LPS: Low protein diet content or LP progeny submitted to the standard environment
MC: Mineralocorticoid
MR: Mineralocorticoid receptor
NP: Regular protein diet content or NP progeny
NPC: Neuron progenitor cells
NPE: Regular protein diet content or NP progeny submitted to the enriched environment
NPS: Regular protein diet content or NP progeny submitted to the standard environment
NPV: Paraventricular nuclei
POMC: Proopiomelanocortin
PVN: Paraventricular nucleus
SGZ: Subgranular zone
SGZ: Subgranular Zone
SNC: Central nervous system
SVZ: Subventricular zone
SVZ: Subventricular Zone
Two-way ANOVA: two-way analysis of variance
VH: Ventral hippocampus

